# Identifying Mangrove-Coral Habitats in the Florida Keys

**DOI:** 10.1101/2020.05.27.119727

**Authors:** Christina A. Kellogg, Ryan P. Moyer, Mary Jacobsen, Kimberly K. Yates

**Author notes:** Corresponding Author: Christina Kellogg, 600 Fourth Street South, St. Petersburg, FL, 33701, USA.

## Abstract

Coral reefs are degrading due to many synergistic stressors. Recently there have been a number of global reports of corals occupying mangrove habitats that provide a supportive environment or refugium for corals, sheltering them by reducing stressors such as oxidative light stress and low pH. This study used satellite imagery and manual ground-truthing surveys to search for mangrove-coral habitats in the Florida Keys and then collected basic environmental parameters (temperature, salinity, dissolved oxygen, pH_NBS_, turbidity) at identified sites using a multi-parameter water quality sonde. Two kinds of mangrove-coral habitats were found in both the Upper and Lower Florida Keys: (1) prop-root corals, where coral colonies were growing directly on (and around) mangrove prop roots, and (2) channel corals, where coral colonies were growing in mangrove channels under the shade of the mangrove canopy, at deeper depths and not in as close proximity to the mangroves. Coral species found growing on and directly adjacent to prop roots included *Porites porites* (multiple morphs), *Siderastrea radians* and *Favia fragum*. Channel coral habitats predominantly hosted *S. radians* and a few *S. siderea*, although single colonies of *Solenastrea bournoni* and *Stephanocoenia intersepta* were observed. Circumstantial evidence suggests additional coral communities existed on mangrove shorelines of oceanside and backcountry islands until destroyed, likely by Hurricane Irma. These mangrove-coral habitats may be climate refugia for corals and could be included in ecosystem management plans and considered for their applications in coral restoration, for example, as a source of adapted genetic resources, places to support growth and acclimation of coral outplants, or natural laboratories to test survival of different genotypes.

## Introduction

Coral reef ecosystems support up to 25% of fisheries in tropical regions and developing nations (Garcia & de Leiva Moreno 2003) and economic and recreational services for more than 100 countries (Burke et al. 2011). Reef framework and shallow, non-coral-dominated habitats serve as natural barriers that protect shoreline ecosystems and coastal communities by reducing hazards from waves, storm surges, and tsunamis for more than 200 million people around the world (Ferrario et al. 2014; Sheppard et al. 2005). However, coral reefs worldwide, and the important ecosystem services they provide, are in a state of critical decline due to a number of synergistic local and global stressors, including coral bleaching, disease, coastal development, overfishing, and nutrient enrichment (Glynn 1984; Precht et al. 2016; Vega Thurber et al. 2013; Weil & Rogers 2011; Yates et al. 2017; Zaneveld et al. 2016).

Coral reef degradation and the causes have been documented since the 1970s (Bruckner & Hill 2009; Gardner et al. 2003; Hughes 1994; Pandolfi et al. 2003), however models suggest nearly 66 % of coral reefs worldwide will continue to undergo rapid degradation over the next few decades due to warming and ocean acidification (Frieler et al. 2013). Ocean acidification results from increasing storage of atmospheric carbon dioxide in the surface ocean, lowering the aragonite saturation state and reducing seawater pH. Coastal acidification caused by eutrophication, coastal upwelling and freshwater inflow also reduces seawater pH and aragonite saturation state. Both of these processes can slow coral growth and contribute to chemical dissolution of reefs (Comeau et al. 2014; Eyre et al. 2018). Reefs in the Florida Keys are already being affected by coastal acidification, likely driven by nutrient inputs resulting in seasonal dissolution of carbonate sediments (Muehllehner et al. 2016) that may be accounting for approximately 15% of seafloor elevation loss in the Upper Florida Keys (Yates et al. 2017). Solar radiation and high water temperatures cause coral bleaching that has resulted in extensive coral mortality as well as predisposing the survivors to coral disease (Miller et al. 2009; Muller et al. 2008; Rogers et al. 2009; Williams & Bunkley-Williams 1990). Coral diseases continue to emerge, including Stony Coral Tissue Loss Disease (SCTLD) which has severely impacted the Florida reef tract since 2014 and is now spreading to the wider Caribbean basin (Precht et al. 2016; Walton et al. 2018; Weil et al. 2019).

Evidence that repeated coral bleaching events (Baker et al. 2008; Eakin et al. 2010; Lesser 2011), coastal and ocean acidification (Fabricius et al. 2011; Kleypas & Yates 2009; Kroeker et al. 2013; Muehllehner et al. 2016; Silverman et al. 2009), coupled with severe and pervasive outbreaks of coral disease will severely impede coral growth within the next few decades (Burke et al. 2011; Hoegh-Guldberg et al. 2007; van Hooidonk et al. 2014) has prompted an urgent search for coral reef systems that provide natural refugia from climate threats. Keppel et al. (2012) define refugia as “habitats that components of biodiversity retreat to, persist in, and can potentially expand from under changing environmental conditions.” The complex interplay among climate, oceanographic, and biological factors that influences susceptibility and resilience of reefs has made identification and characterization of such refugia for corals challenging.

Conservation and management strategies include the establishment of marine protected areas with environmental conditions that promote coral resiliency. Focus has been placed on identifying reefs with low exposure to or potential for adaptation to climate threats, and reduced local anthropogenic impacts (Keller et al. 2009; Mumby & Steneck 2008; Salm et al. 2006; West & Salm 2003). Recent studies have identified only one reef in the Florida Keys as a potential refuge from ocean acidification (Manzello et al. 2012). Mangrove communities, while often near coral reef ecosystems, are not typically thought of as having suitable conditions for coral recruitment and growth due to high sedimentation rates, lack of suitable substratum, and inadequate water quality. Further, ecological surveys of Florida mangroves from the 1930s and 1980s made no mention of the presence of corals when detailing associated fauna (Davis 1940; Odum et al. 1982). However, a number of recent studies have identified several locations around the world with corals growing on or near mangrove prop roots (Camp et al. 2019; Camp et al. 2017; Macintyre et al. 2000; Rogers 2009; Rogers 2017). In some of these habitats, mangroves are sheltering corals even in the face of extreme variability in pH, dissolved oxygen, and temperature, resulting in lower incidences of bleaching and high rates of recovery (Camp et al. 2019; Camp et al. 2017; Yates et al. 2014). The mangrove-canopy shading reduces light stress and a combination of hydrodynamic and biogeochemical processes in some of these mangrove-coral habitats can locally buffer pH (Yates et al. 2014). This is the first study to systematically search for and identify mangrove-coral habitats in the Florida Keys and provide a basic environmental characterization of them.

## Materials & Methods

### Site selection

Several areas in the Upper and Lower Florida Keys were identified as target areas based on previous unpublished observations by the authors, and/or anecdotal personal communication from other researchers that have worked in the Florida Keys, that corals had been previously observed in or near mangrove shorelines. Additional target areas were chosen by using satellite images from Google Earth Pro (Version 7.3, Google LLC, Mountain View CA, USA) to identify mangrove shorelines that were adjacent to tidal channels with one or more of the following criteria: (i) deep enough to support corals at all stages of the tidal cycle, (ii) deep enough, or with visible evidence (e.g., tidal deltas present) to suggest strong current flow, (iii) clear water, (iv) a connection to the open ocean, and (v) areas where hard substrate was mapped adjacent to mangroves on the Florida Fish and Wildlife Conservation Commission (FWC)’s Unified Reef Map (https://myfwc.com/research/gis/regional-projects/unified-reef-map/). Some mangrove-lined channels that could not be easily observed via satellite were included for ground truthing. Heavily built areas (e.g., Key Largo, Marathon, Key West) were avoided since they were likely to have fewer mangrove-lined shorelines and poorer water quality.

### Field surveys

Maps of target areas were used to guide visual surveys of mangrove shorelines and channels. Surveys were conducted between 0800–1700 for optimal lighting. Areas in the Upper Keys (Biscayne Bay/Card Sound/Largo Sound) were surveyed 4–8 October 2019 and areas in the Lower Keys (between Big Pine Key and Boca Chica Key) were surveyed 7–11 January 2020. Depending on accessibility, surveys for the presence of corals growing on mangrove prop roots or in channels shaded by the mangrove canopy were conducted by boating at very low speed, paddleboard, or snorkeling. Areas surveyed were recorded using a hand-held wide-area-augmentation-system (WAAS)-corrected global position system (GPS). When corals were located in mangrove habitats, each coral species and their corresponding abundances were recorded and representative photographs of the corals were taken. The following environmental parameters were measured using a hand-held multi-parameter water-quality sonde (YSI ProDSS, Xylem Inc., Yellow Springs OH, USA): water temperature (degrees Celsius), salinity, dissolved oxygen (mg/L), turbidity (Formazin nephelometric units, FNU), pH_NBS_ (to estimate relative differences in pH between mangrove-coral and reference habitats), and pressure (dbar) to estimate water depth (meters).

### Area surveyed

Way points from the GPS were plotted daily after each survey in Google Earth Pro. The Google Earth KMZ file was then imported into ArcGIS Pro (Esri Inc., Redlands CA, USA) to create maps with track lines to represent the surveyed areas. The length of the track lines was calculated by ArcGIS Pro based on the WGS84 Web Mercator (Auxiliary Sphere) projection used for the National Agriculture Imagery Program (NAIP) base map. The calculated length of the track lines was summed to obtain the estimated kilometers of mangrove shoreline surveyed.

## Results

The total linear distance of mangrove shoreline that was surveyed during this project was approximately 76 km. The surveys identified two kinds of mangrove-coral habitats in the Florida Keys: (1) prop-root corals, where colonies were growing directly on (and in close proximity to, defined as less than 0.5 m) mangrove prop roots, and (2) channel corals, where colonies were growing in tidal channels between mangrove shorelines, such that the corals were shaded during at least part of the day by the mangrove canopy, but not close to prop roots.

### Upper Keys Surveys

Approximately 55 km of mangrove shoreline in the Upper Florida Keys, including parts of Card Sound and Largo Sound, were surveyed 4–8 October 2019 (Figs. 1 and 2). An additional mangrove-lined tidal channel (not shown in Fig. 1) was surveyed in North Key Largo from the southern end of Card Sound to an impassable bridge clearance beneath Card Sound Road. In this channel, the water was very turbid, appearing opaque dark brown in color, and no corals were observed there. Both prop-root and channel coral habitats were observed in the Upper Keys and environmental data were collected at representative sites (Table 1). All sites with prop-root corals were found along the northern side of a deeply incised channel next to Swan Key (inset, Fig. 1) and featured at least two different morphotypes of *Porites porites* and one encrusting *Siderastrea radians* colony (Table 1, Fig. 3). Prop-root corals in the Upper Keys ranged in size (longest nominal axis) from 2–20 cm. Channel coral habitat was found in mangrove-lined tidal channels cutting through the interior of islands (e.g., Swan Creek, inset, Fig. 1), and featured small colonies of *Siderastrea siderea, S. radians*, and *Stephanocoenia intersepta* (Table 1, Fig. 3). Clusters of small coral colonies were occasionally observed in some wider interior channels that were not being shaded by mangroves (Angelfish Key, Old Rhodes Key). Channel corals in the Upper Keys ranged in size (longest nominal axis) from 2–25 cm. All of the tidal channels surveyed around Largo Sound (Fig. 2) had discolored water with high turbidity and low visibility, and no corals were seen in spite of previously reported anecdotal sightings. A location in these channels was chosen to collect environmental data as a non-coral-habitat reference site for comparison (Table 1, Fig. 2).

**Table 1:** Mangrove-coral habitat data for Upper Florida Keys sites. Sites indicate locations of channel corals (C), prop-root corals (P) or no corals (NC) as depicted in Figures 1 and 2. Brackets contain the number of coral colonies observed per species at a given site.

| Date | Local Time (EDT) | Site | Location | Habitat | Coral Sp. | Lat/Long | Temp (°C) | Salinity | DO (mg/L) | pH<sub>NBS</sub> | Turbidity (FNU) | Tidal State | Depth (m) |
| --- | --- | --- | --- | --- | --- | --- | --- | --- | --- | --- | --- | --- | --- |
| 10/5/19 | 10:00 | C1 | Swan Creek | channel | *Siderastrea siderea* [3],<br>*Siderastrea radians* [8],<br>*Stephanoeonia intersepta* [1] | 25.34798<br>- 80.24977 | 27.8 | 36.32 | 4.47 | 7.93 | -0.9 | falling | 1.2 |
| 10/5/19 | 10:15 | C2 | Swan Creek | channel | *S. siderea* [2],<br>*S. radians* [2] | 25.34793<br>-80.24963 | 28.0 | 36.33 | 4.65 | 7.93 | -1.1 | falling | 1.4 |
| 10/5/19 | 10:25 | C3 | Swan Creek | channel | *S. radians* [1] | 25.34819<br>- 80.24959 | 28.0 | 36.34 | 4.80 | 7.91 | -1.1 | falling | 1.37 |
| 10/5/19 | 12:30 | C4 | Angelfish Creek | channel | *S. radians* [56],<br>*Solenastrea bournoni* [1] | 25.3326<br>-80.2645 | 28.2 | 36.57 | 4.25 | 7.78 | -0.3 | rising | 1.96 |
| 10/6/19 | 10:45 | P1 | Swan Key | prop root | *Porites porites* [1]<br>*S. radians* [1] | 25.34598<br>- 80.24873 | 27.6 | 36.86 | 5.35 | 7.96 | -0.8 | falling | 0.39 |
| 10/6/19 | 11:40 | P2 | Swan Key | prop root | *P. porites* [1] | 25.34755<br>- 80.25092 | 27.8 | 36.87 | 5.76 | 7.99 | -0.9 | falling | 0.48 |
| 10/6/19 | 12:05 | P3 | Swan Key | prop root | *P. porites* [1] | 25.34757<br>- 80.24982 | 27.9 | 36.78 | 6.03 | 8.00 | -1.0 | falling, nearly slack | 0.46 |
| 10/7/19 | 13:34 | NC | Key Largo Negative control | channel | N/A | 25.1584<br>-80.3783 | 27.5 | 36.77 | 2.27 | 7.24 | -0.5 | rising | 0.95 |

Abbreviations: EDT – Eastern Daylight Time (GMT -4), Lat/Long – latitude and longitude in decimal degrees, DO – (optical) dissolved oxygen, FNU – Formazin Nephelometric Units, N/A – not applicable.

**Figure 1.**
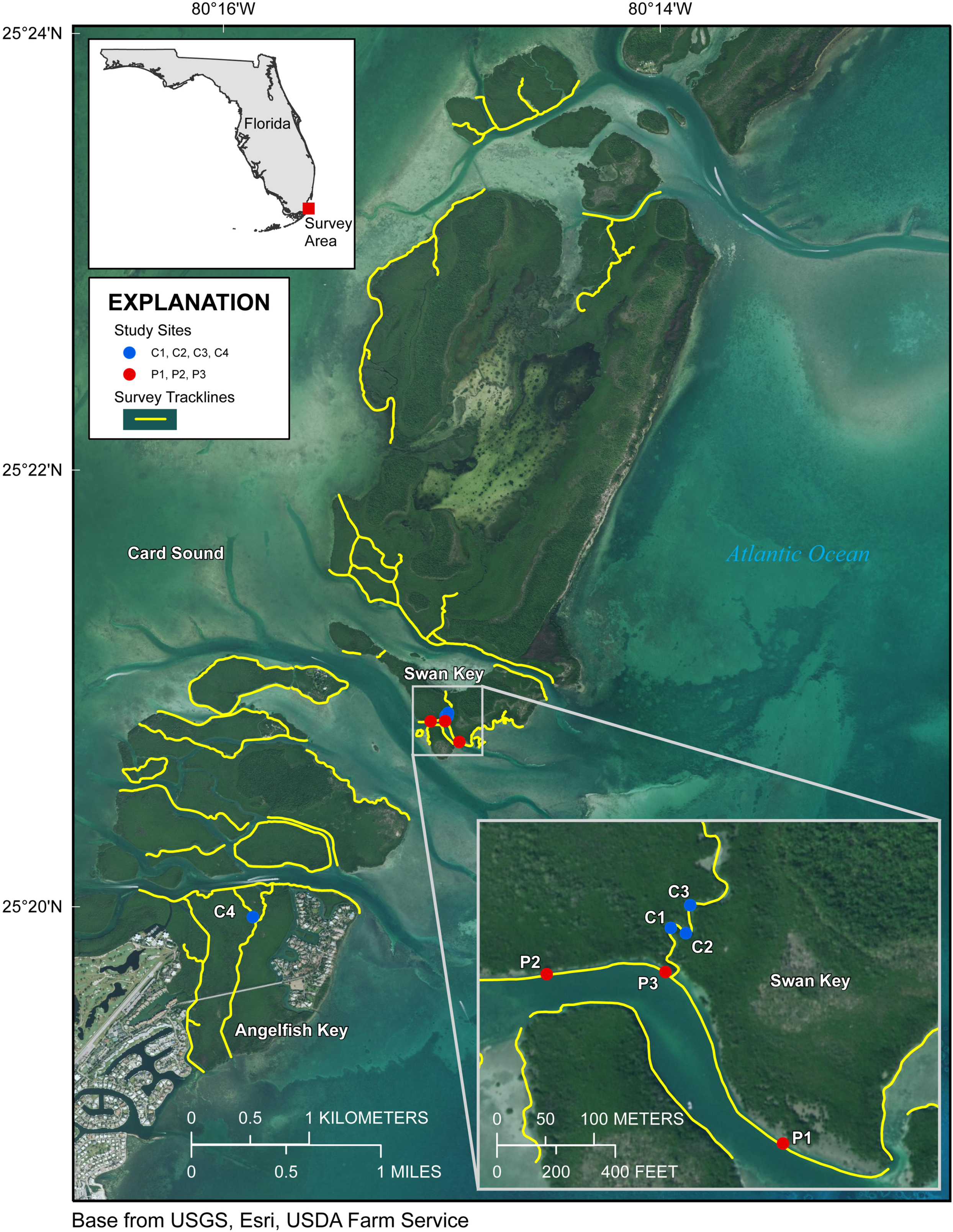
Upper Florida Keys surveys in the vicinity of Card Sound. Yellow lines indicate shoreline and channels surveyed. Red points labeled P1, P2, and P3 indicate prop-root-coral sites described in Table 1. Blue points labeled C1, C2, C3 and C4 indicate channel-coral sites described in Table 1. Map image is the intellectual property of Esri and is used herein under license. Copyright ©2019 Esri and its licensors. All rights reserved.

**Figure 2.**
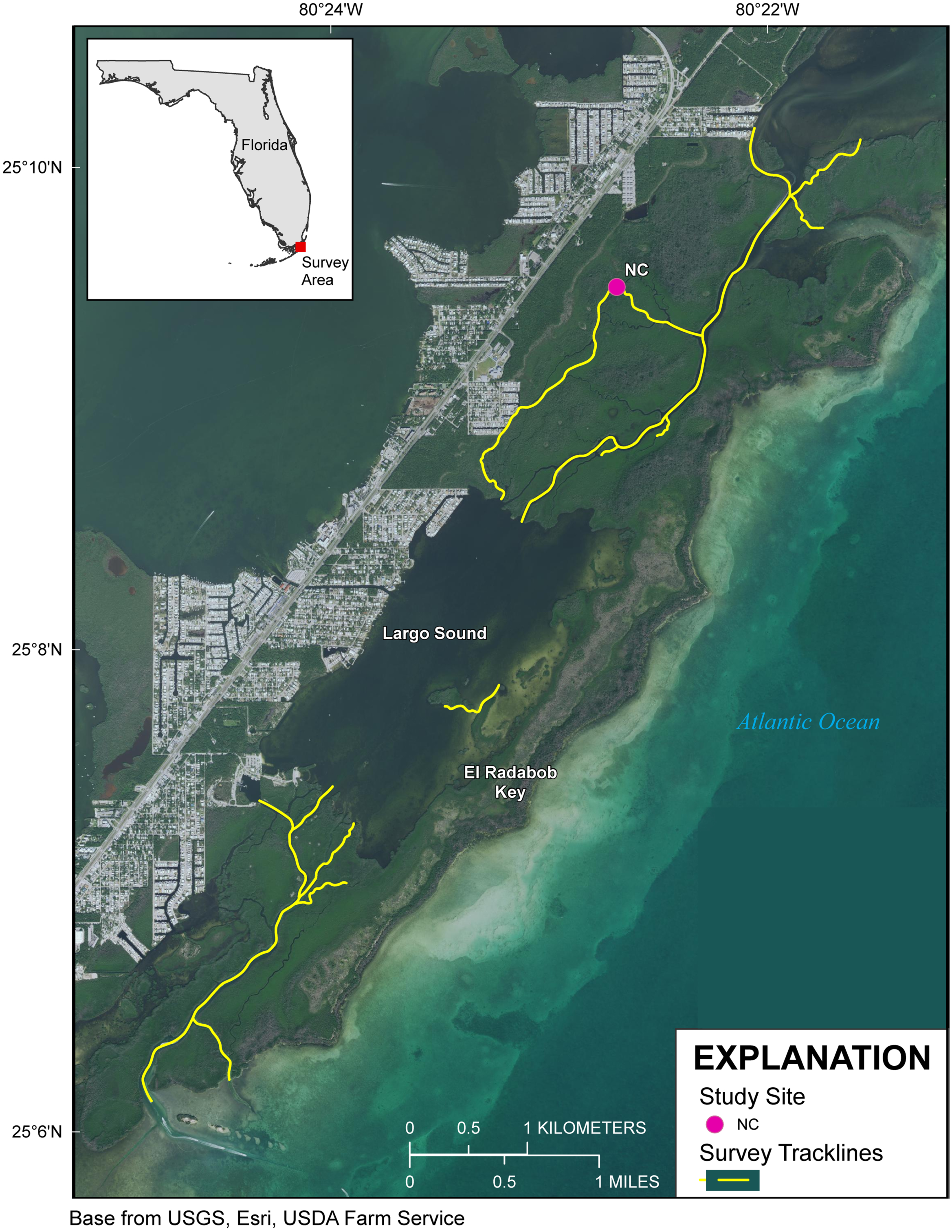
Upper Florida Keys surveys around Largo Sound. Yellow lines indicate shoreline and channels surveyed. Purple point labeled NC indicates reference site sampled for environmental parameters described in Table 1. Map image is the intellectual property of Esri and is used herein under license. Copyright ©2019 Esri and its licensors. All rights reserved.

**Figure 3.**
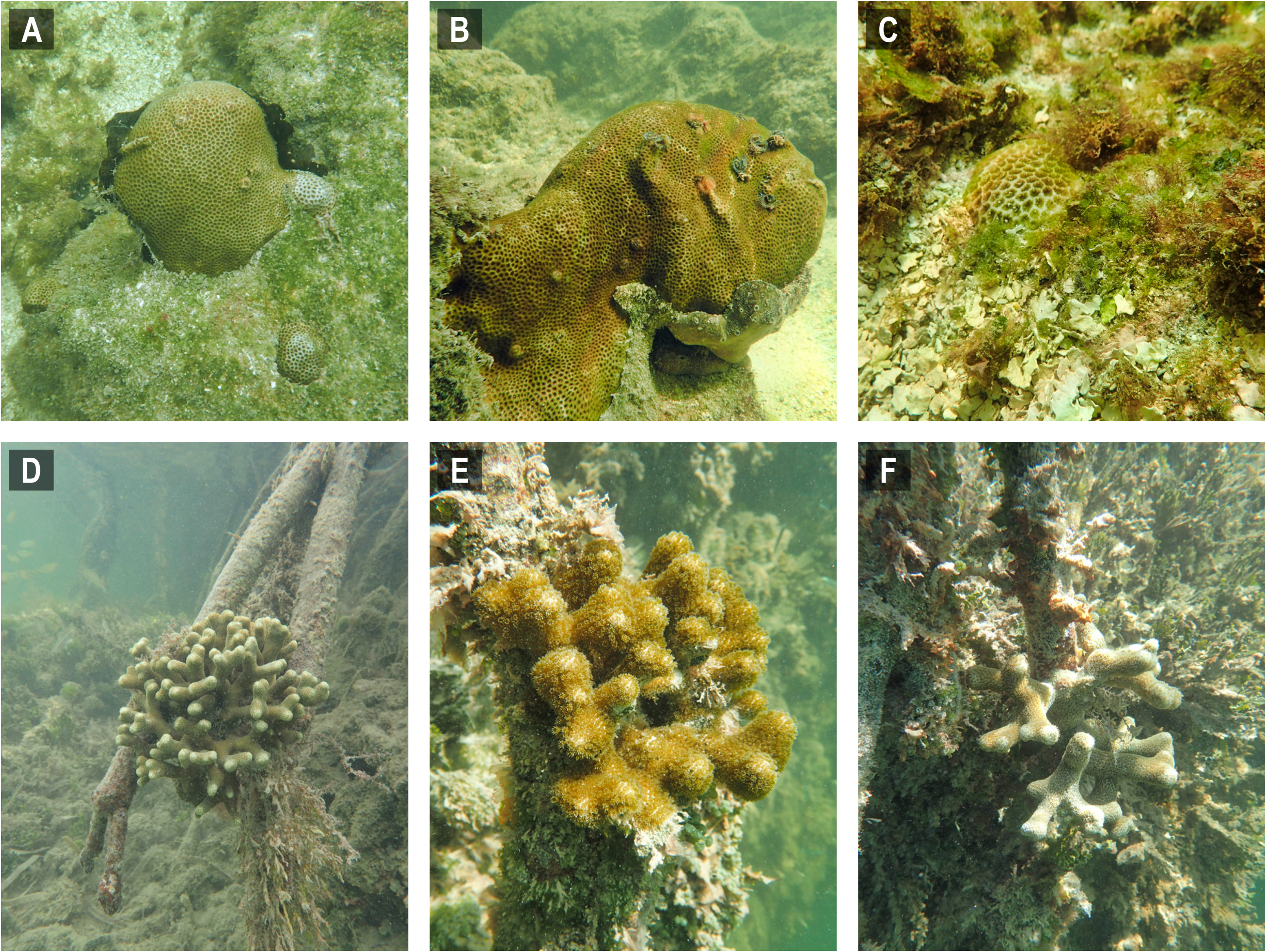
Selected images of mangrove-coral habitats in the Upper Florida Keys. Panel A = *Siderastrea radians*, site C1, B = *S. radians*, site C1, C = *S. radians*, site C2, D = *Porites porites*, site P1, E = *P. porites*, site P2, F = *P. porites*, site P3.

A two-sample t-test assuming unequal variances and two tails was performed in Microsoft Excel (Microsoft Inc., Redmond WA, USA) to test for differences between Upper Keys channel coral habitats and prop-root-coral habitats based on the data in Table 1. There were no significant differences in temperature, pH, or turbidity between the two habitat types. However, there were significant differences in salinity (channel corals mean 36.39 ± 0.014; prop-root corals mean 36.84 ± 0.002; t_stat_ = −6.71, d.f. = 4, p = 0.003) and dissolved oxygen (channel corals mean 4.54 ± 0.056; prop-root corals mean 5.71 ± 0.117; t_stat_ = −5.08, d.f. = 3, p = 0.015). This may reflect the difference between the physical characteristics (depth, current velocity, and oceanic influence) on the channel with prop-root corals versus the tidal creek hosting corals mid-channel (Fig. 1). Because environmental data were only collected at one non-coral reference site (Fig. 2), it is not possible to test for significant differences between target and reference habitats; however, both dissolved oxygen concentrations and pH values were much lower at the Upper Keys reference site compared to both types of mangrove-coral habitats (Table 1).

### Lower Keys Surveys

Approximately 21 km of linear mangrove shoreline was surveyed in the Lower Florida Keys between Big Pine Key and Boca Chica Key from 7–11 January 2020 (Fig. 4). Both prop-root- and channel-coral habitats were observed in the Lower Keys and environmental data were collected at representative sites (Tables 2 and 3). Although surveys included mangrove shorelines on the ocean side islands and in the more protected backcountry islands, all prop-root-coral sites were found in natural tidal channels or man-made canals connecting the Atlantic Ocean with Upper Sugarloaf Sound, with the exception of Park Channel, which connects Lower and Upper Sugarloaf Sounds (Fig. 4). The most common species observed growing on prop roots was *Porites porites* (Table 2, Fig. 5). The highest diversity and largest abundance of individual colonies of prop-root and shaded corals (species: *Porites porites, Siderastrea radians*, and *Favia fragum*) was found in a man-made canal dredged through Pleistocene bedrock (Miami Limestone formation) of Sugarloaf Key, connecting Upper Sugarloaf Sound and the Atlantic Ocean. This dredged canal runs parallel to Sugarloaf Boulevard and passes under the Loop Road Bridge. It is cataloged in the Monroe County Canal Management Master Plan as “430 Sugarloaf Key Merged Canal.” Channel corals, mainly *S. radians*, were observed in Tarpon Creek and throughout the length of 430 Sugarloaf Key Merged Canal (Table 2, Fig. 5). Prop-root corals in the Lower Keys ranged in size (longest nominal axis) from 5–25 cm and channel corals ranged between 1–35 cm in size.

**Table 2:** Mangrove-coral habitat data for Lower Florida Keys sites. Sites indicate locations of channel corals (C) and prop-root corals (P) as depicted in Figure 4. Shaded cells indicate revisits to a site at a different date/time. Brackets contain the number of coral colonies observed per species at a given site.

| Date | Local Time (EDT) | Site | Location | Habitat | Coral Sp. | Lat/Long | Temp (°C) | Salinity | DO (mg/L) | pH<sub>NBS</sub> | Turbidity (FNU) | Tidal State | Depth (m) |
| --- | --- | --- | --- | --- | --- | --- | --- | --- | --- | --- | --- | --- | --- |
| 1/7/20 | 08:40 | C5 | Tarpon Creek | channel | *Siderastrea radians* [>10] | 24.628111<br>-81.51174 | 19.9 | 36.28 | 4.12 | 8.09 | 0.5 | slack | 0.25 |
| 1/7/20 | 16:45 | P4 | Tarpon Canal | prop root | *Porites porites* [1] | 24.631081<br>-81.512361 | 22.0 | 36.34 | 9.20 | 8.41 | 1.9 | rising | 0.32 |
| 1/11/20 | 10:05 | P4 | Tarpon Canal | prop root | *P. porites* [1] | 24.631081<br>-81.512361 | 22.8 | 36.66 | 5.00 | 8.07 | 1.3 | rising | 0.25 |
| 1/9/20 | 09:28 | P5 | Park Channel | prop root | *P. porites* [1] | 24.647889<br>-81.558723 | 20.1 | 36.24 | 6.45 | 8.33 | -0.7 | falling | 0.06 |
| 1/9/20 | 09:40 | P6 | Park Channel | prop root | *S. radians* [1] | 24.647813<br>-81.558667 | 20.1 | 36.27 | 6.61 | 8.35 | -0.8 | falling | 0.03 |
| 1/9/20 | 11:26 | C6 | 430 Sugarloaf Key Merged Canal | channel | *S. radians* [7]. | 24.618799.<br>-81.531484 | 20.1 | 36.39 | 6.62 | 8.4 | 0.2 | falling | 0.53 |
| 1/9/20 | 13:42 | P7 | 430 Sugarloaf Key Merged Canal | prop root | *P. porites* [4],<br>*S. radians* [4] | 24.630163<br>-81.541481 | 21.5 | 36.37 | 8.43 | 8.70 | 0.3 | falling,<br>almost slack | 0.08 |
| 1/11/20 | 11:46 | P7 | 430 Sugarloaf Key Merged Canal | prop root | *P. porites* [1] | 24.630163<br>-81.541481 | 23.5 | 36.46 | 6.99 | 8.38 | 1.2 | falling | 0.15 |
| 1/11/20 | 14:03 | P7 | 430 Sugarloaf Key | prop root | *P. porites* [1] | 24.630163<br>-81.541481 | 24.3 | 36.45 | 7.88 | 8.50 | 5.10 | Falling | 0.08 |

|  |  |  | Merged Canal |  |  |  |  |  |  |  |  |  |  |
| --- | --- | --- | --- | --- | --- | --- | --- | --- | --- | --- | --- | --- | --- |
| 1/11/20 | 11:47 | P8 | 430 Sugarloaf Key Merged Canal | prop root | *Favia fragum* [1] | 24.630251<br>-81.541658 | 23.4 | 36.47 | 6.84 | 8.38 | 0.7 | falling | 0.049 |
| 1/9/20 | 13:00 | P9 | 430 Sugarloaf Key Merged Canal | prop root | *P. porites* [1] | 24.630322<br>-81.541749 | ND | ND | ND | ND | ND | ND | ND |
| 1/9/20 | 13:03 | P10 | 430 Sugarloaf Key Merged Canal | prop root | *P. porites* [1] | 24.630332<br>-81.541804 | ND | ND | ND | ND | ND | ND | ND |
| 1/9/20 | 13:10 | P11 | 430 Sugarloaf Key Merged Canal | prop root | *P. porites* [1] | 24.630486<br>-81.541895 | ND | ND | ND | ND | ND | ND | ND |
| 1/9/20 | 13:15 | P12 | 430 Sugarloaf Key Merged Canal | prop root | *P. porites* [1] | 24.630723<br>-81.542168 | ND | ND | ND | ND | ND | ND | ND |
| 1/9/20 | 13:20 | P13 | 430 Sugarloaf Key Merged Canal | prop root | *P. porites* [1] | 24.630775<br>-81.542177 | ND | ND | ND | ND | ND | ND | ND |
| 1/9/20 | 13:25 | P14 | 430 Sugarloaf Key Merged Canal | prop root | *P. porites* [1] | 24.631218<br>-81.542646 | ND | ND | ND | ND | ND | ND | ND |
| 1/9/20 | 13:30 | P15 | 430<br>Sugarloaf<br>Key<br>Merged<br>Canal | prop root | *P. porites* [1] | 24.631868<br>-81.543327 | ND | ND | ND | ND | ND | ND | ND |
| 1/9/20 | 16:25 | P16 | Five Mile<br>Creek | prop root | *P. porites* [1] | 24.649801<br>-81.596691 | 20.7 | 36.24 | 7.62 | 8.60 | -0.8 | rising | 0.15 |

Abbreviations: EDT – Eastern Daylight Time (GMT -4), Lat/Long – latitude and longitude in decimal degrees, DO – (optical) dissolved oxygen, FNU – Formazin Nephelometric Units, ND – not determined.

**Table 3:** Comparison between environmental parameters in prop-root coral habitats and non-target habitats in the Lower Florida Keys. All data collected on January 11, 2020. Shading indicates prop-root coral habitats.

| Local Time (EDT) | Habitat | Description | Lat/Long | Temp (°C) | Salinity | DO (mg/L) | pH<sub>NBS</sub> | Turbidity (FNU) | Tidal State | Depth (m) | Notes |
| --- | --- | --- | --- | --- | --- | --- | --- | --- | --- | --- | --- |
| 09:50 | Coastal Atlantic | Oceanside, in boat channel outside of Tarpon Canal | 24.6305<br>-81.5066 | 22.6 | 36.37 | 6.57 | 8.30 | 3.7 | rising | 0.20 | Open water, no proximity to corals or mangroves |
| 09:55 | Mangrove canal | Oceanside entrance, Tarpon Canal | 24.6316<br>-81.5091 | 22.8 | 36.52 | 6.40 | 8.24 | 4.4 | rising | 0.01 | Mid-channel, mangroves line channel edges |
| 10:05 | Prop-root coral (P4) | Interior, Tarpon Canal | 24.6311<br>-81.5124 | 22.8 | 36.66 | 5.00 | 8.07 | 1.3 | rising | 0.25 | Single mature *Porites* colony growing on mangrove prop root, north side of canal |
| 10:06 | Canal-coral proximity | Interior, Tarpon Canal | 24.6311<br>-81.5124 | 22.8 | 36.59 | 5.57 | 8.16 | 1.5 | rising | 0.22 | Mid-channel reading parallel with site Prop-root coral site P4 |
| 10:16 | Mangrove canal | Interior, Tarpon Canal | 24.6307<br>-81.5146 | 23.0 | 36.70 | 5.44 | 8.10 | 1.5 | rising | 0.06 | Against mangroves without corals, north side of canal almost opposite opening to Tarpon Creek |
| 10:32 | Inland waterway | Upper Sugarloaf Sound | 24.6382<br>-81.5276 | 23.1 | 36.42 | 6.91 | 8.48 | 3.0 | slack | 0.88 | Open water, mid-basin, no proximity to corals or mangroves |
| 11:44 | Channel coral & prop-root coral area | 430 Sugarloaf Key Merged Canal | 24.6302<br>-81.5415 | 23.5 | 36.42 | 7.10 | 8.38 | 0.8 | falling | 0.089 | Mid-channel reading parallel with prop-root coral site P7 |
| 11:46 | Prop-root coral (P7) | 430 Sugarloaf Key Merged Canal | 24.6302<br>-81.5415 | 23.5 | 36.46 | 6.99 | 8.38 | 1.2 | falling | 0.15 | Multiple *Porites* colonies growing on mangrove prop roots of same plant, east side of canal |
| 11:47 | Prop-root coral (P8) | 430 Sugarloaf Key Merged Canal | 24.6303<br>-81.5417 | 23.4 | 36.47 | 6.84 | 8.38 | 0.7 | falling | 0.049 | Small colony of *Favia* growing on mangrove prop root, west side of canal |
| 11:53 | Mangrove canal | 430 Sugarloaf Key Merged Canal | 24.6320<br>-81.5433 | 23.8 | 36.39 | 7.48 | 8.43 | 1.1 | falling | 0.024 | Mid-channel, near marker pole at Sugarloaf Sound entrance to canal |
| 14:52 | Channel coral area | 430 Sugarloaf Key Merged Canal | 24.6205<br>-81.5319 | 24.3 | 36.43 | 7.69 | 8.67 | 3.20 | falling | 0.08 | Mid-channel, thick mangroves along sides, channel corals on rock walls but no prop-root corals |

Abbreviations: EDT – Eastern Daylight Time (GMT -4), Lat/Long – latitude and longitude in decimal degrees, DO – (optical) dissolved oxygen, FNU – Formazin Nephelometric Units.

**Figure 4.**
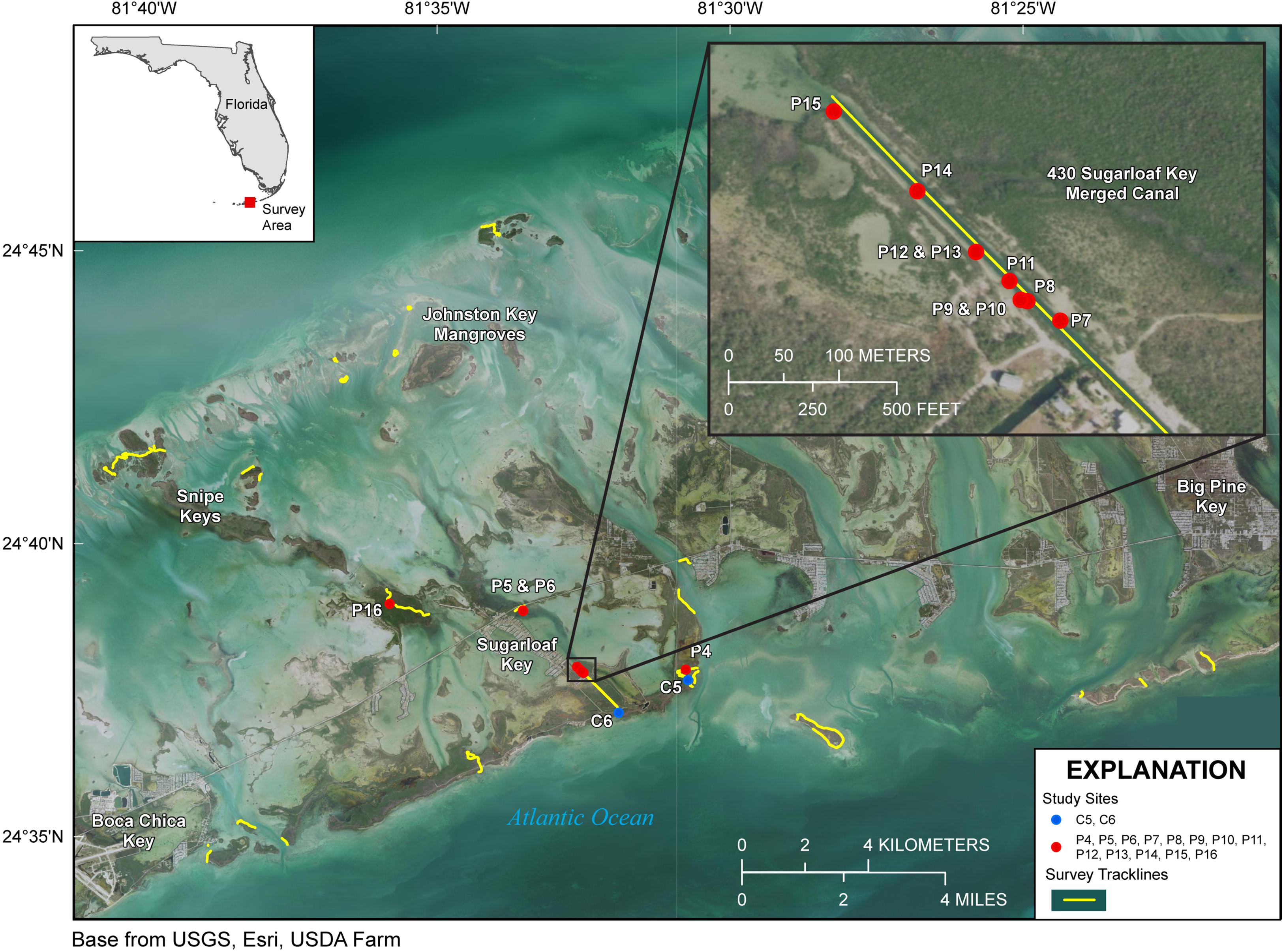
Lower Florida Keys Surveys. Yellow lines indicate shoreline and channels surveyed. Red points labeled P4 to P16 indicate prop-root-coral sites described in Table 2. Blue points labeled C5 and C6 indicate channel-coral sites described in Table 2. Map image is the intellectual property of Esri and is used herein under license. Copyright ©2019 Esri and its licensors. All rights reserved.

**Figure 5.**
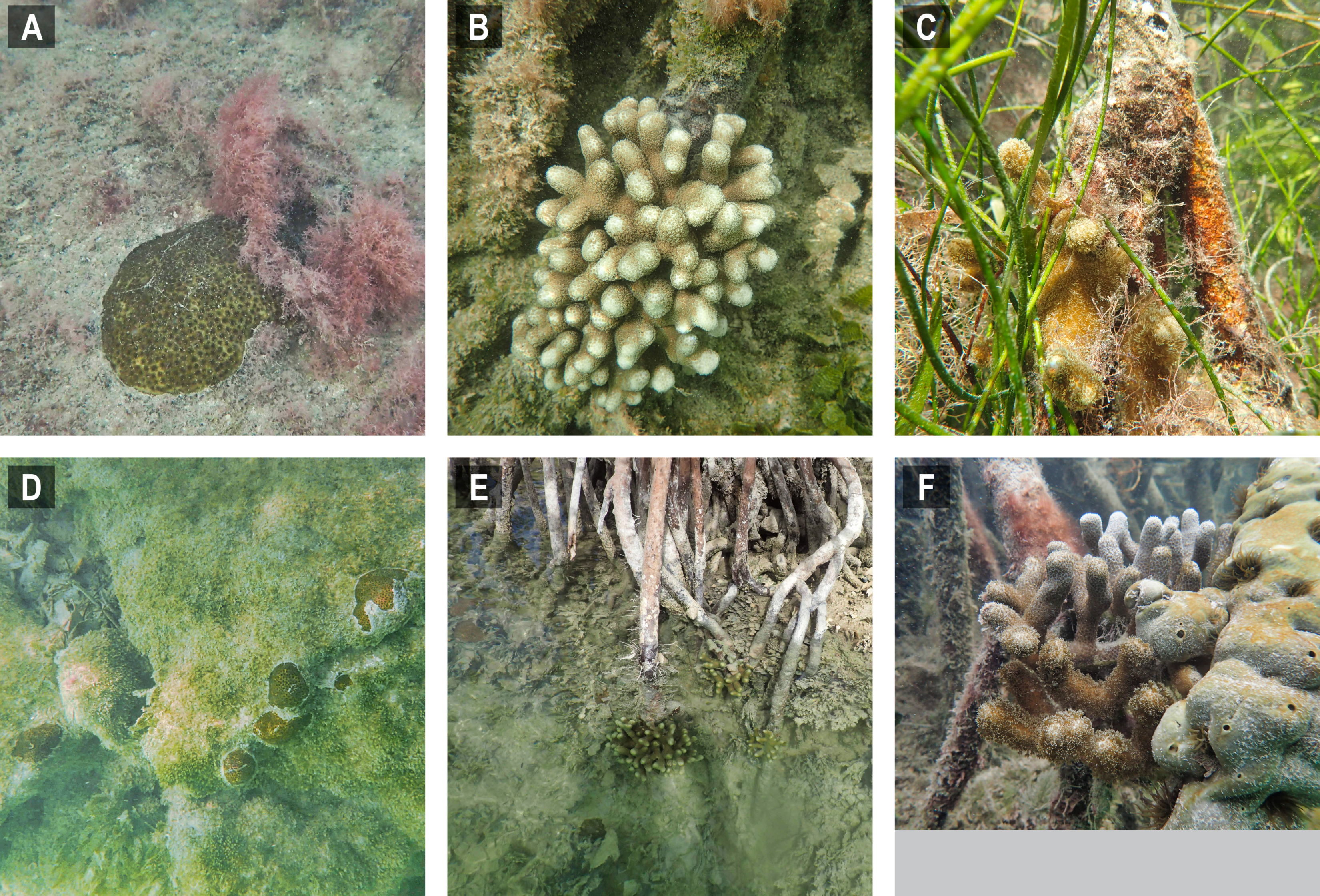
Selected images of mangrove-coral habitats in the Lower Florida Keys. Panel A = *Siderastrea radians*, site C5, B = *Porites porites*, site P4, C = *P. porites*, site P5, D = *S. radians*, site C6, E = *P. porites*, site P7, F = *P. porites*, site P16.

A two-sample t-test assuming unequal variances and two tails was performed in Microsoft Excel to test for differences between Lower Keys channel-coral habitats and prop-root-coral habitats based on the data in Table 2. The only environmental variable that was significantly different was temperature and that can be attributed to the differences between days and sampling times, whereby more prop-root-coral sites were visited in the afternoon or were visited on 11 January, when surface-water temperatures were above 22°C. Both prop-root- and channel-coral habitats in the Lower Keys occurred in inland tidal channels and canals, so it is not unexpected that major differences were not detected among measured environmental parameters at each type of site.

A two-sample t-test assuming unequal variances and two tails was performed in Microsoft Excel to test for differences between the prop-root-coral sites and reference sites based on data in Table 3 that were collected on the same day to minimize astochastic variability introduced by sampling at different times of day on multiple days. Reference sites were those with open water, mangrove-lined shorelines or confined tidal channels that did not serve as habitat for prop-root or channel corals. The only environmental parameter in the Lower Keys that was significantly different between prop-root-coral sites and reference sites was turbidity (Table 3; prop-root-coral sites mean 1.1 ± 0.115; reference sites mean 2.8 ± 1.630; t_stat_ = 3.16, d.f. = 6, p = 0.02).

Note that there was no combined analysis of environmental data from Upper and Lower Keys surveys because they were conducted during different seasons. Seasonality would not affect the presence or absence of corals, but could confound any observed differences in environmental data.

## Discussion

The mangrove-coral habitats identified in the Florida Keys during this project did not appear to be subject to extreme environmental variability like those reported in New Caledonia and Australia (Camp et al. 2019; Camp et al. 2017). In that respect, the sites we surveyed were more similar to the mangrove-coral habitats of the U.S. Virgin Islands (Rogers 2017; Yates et al. 2014); however, the Keys habitats hosted substantially lower diversity of corals and were dominated by stress-resilient species, primarily *P. porites* and *S. radians* (Lirman et al. 2002). However, we did document the presence of other coral species, more commonly in the channel-coral habitats than the prop-root-coral habitats (Tables 1 and 2): *Favia fragum, S. siderea, So. bournoni, St. intersepta*. Our environmental measurements (Tables 1-3) fall within the normal mean ranges measured on Florida Keys inshore and offshore reefs in recent decades: temperatures were compared against the multi-decadal data available from National Oceanic and Atmospheric Administration (NOAA)’s National Data Buoy Center (www.ndbc.noaa.gov); salinity and dissolved oxygen were compared against water quality data from the Florida Keys National Marine Sanctuary (Briceño & Boyer 2015). The pH values we measured are relative and therefore could not be compared accurately to absolute data. However, it is likely the extremes in environmental variables rather than the means that determine the ability of corals to survive in these particular locations.

The Florida Keys episodically experience cold fronts that can push water temperatures below the 16–18°C lower limit of tropical scleractinian tolerance for several days causing mass coral mortality, as occurred in 1977 and 2010 (Lirman et al. 2011; Roberts et al. 1982). However, for cold or cool weather pulses of lesser duration, we hypothesize that mangrove-coral habitats may be somewhat thermally buffered by the microclimate effect of the mangrove canopy and the retention of heat by peat and porewaters (Osland et al. 2019).

The full extent of benefits that may be derived by corals in mangrove habitats remains to be determined. The experimentally proven advantages in the Virgin Islands included carbonate system buffering and reduction of oxidative stress via shading (Yates et al. 2014). Other observed benefits include lower incidence of bleaching and/or more rapid recovery from bleaching (Camp et al. 2017; Yates et al. 2014). Given that bleaching has been linked to increased subsequent mortality by disease (Miller et al. 2009; Rogers et al. 2009), these mangrove-coral habitats may also provide indirect protection against coral disease. In the Lower Keys, we detected a significant difference in turbidity between coral and reference habitats. High turbidity in mangrove-adjacent waters is typically caused by the high input of dissolved and particulate organic matter derived from the direct productivity of the mangrove forest (Alongi 2014). Some components of dissolved organic matter can function as antioxidants and this activity has been documented to be particularly high in Florida mangrove environments, likely due to their release of polyphenols and tannins, which are known antioxidants (Romera-Castillo & Jaffé 2015). This may be an added benefit provided to corals by mangroves in addition to the physical shading. It was noted that corals growing on prop roots occurred where the roots reached deep enough under water to not expose the coral at low tide, which, when combined with higher turbidity, also allows for light attenuation and thus less oxidative stress. In the Lower Keys, the majority of prop-root corals were found on the western side of the canals, which was shaded from the afternoon sun by the mangrove canopy. The Tarpon Canal coral (P4) and the three prop-root corals in the Upper Keys (P1-P3) were all on the north side of channels. In 430 Sugarloaf Key Merged Canal we found corals growing on prop roots on both the east (P7) and west (P8-P15) sides of the channel. However, the corals growing on the very shallow substrate directly adjacent to mangroves in 430 Sugarloaf Key Merged Canal were found primarily on the western side of the channel where they were shaded by afternoon sun.

The prop-root corals in the Upper Keys occurred where it was hypothesized they would be, on the edges of deep channels with fast-moving currents that were directly connected to open-ocean water (Fig. 1). However, in the Lower Keys, all the mangrove-coral habitats were observed in protected internal/inland water bodies (Fig. 4) rather than on mangrove islands closer to oceanic water (i.e., along the Atlantic-facing side of offshore islands or along the Gulf of Mexico coast of the backcountry islands). In fact, the most heavily populated area of mangrove-coral habitat (both prop-root and channel corals) surveyed was in the 430 Sugarloaf Key Merged Canal (inset, Fig. 4). This man-made canal runs for a length of 1,840 meters and when surveyed for water quality in 2013, it was noted to have “excellent biodiversity,” possibly referring to the visible coral colonies growing in it (Monroe County Canal Management Master Plan, September 20, 2013). Using spatio-temporal modeling, a recent paper determined that SCTLD appears to move via bottom currents and sediment (Muller et al. 2020), so the disease may not easily transmit into channels and canals where corals are growing, affording them some protection. Further, Bayesian models suggested that corals on high-diversity reefs and on deep reefs were at greater risk of SCTLD than corals on shallow and low-diversity reefs (Muller et al. 2020). Combined, these modeling results indicate that these inland tidal channels and man-made canals may benefit from physical/hydrographic impediments to the movement of the coral disease. It is worth noting that the colony sizes observed growing on prop-roots (Figs. 3 and 5) indicate that these corals were present prior to the stony coral tissue loss disease outbreak moving through these parts of the Florida reef tract in 2016–2018. However, the main corals observed growing on prop roots were *P. porites*, a species which is less susceptible to SCTLD and has been shown by Florida Keys coral surveys to be increasing in abundance in spite of the outbreak (Muller et al. 2020; Walton et al. 2018).

Could these mangrove-coral habitats be functioning as refugia? We argue the possibility exists for these environments to be (i) thermal refugia (via microclimate insulation against cold and shading against heat), (ii) acidification refugia (via buffering pH), (iii) oxidative stress refugia (via shading and mangrove antioxidants), (iv) disease refugia (via hydrographic transmission limitation of the channels), (v) storm refugia (inland tidal creeks and channels may be more protected from heavy wave action and sedimentation), or (vi) various combinations thereof. It is worth testing these hypotheses to determine whether these Florida Keys mangrove-coral habitats could offer specific protection for corals. If so, that might make them suitable as temporary or longer-term nurseries to support growth and acclimation of coral outplants or natural laboratories to test survival of different coral genotypes. Both prop-root and channel corals identified in this study could be sources for genetic alleles adapted to extreme environments, worth investigating for inclusion in restoration efforts seeking to increase genetic diversity (Baums 2008; Baums et al. 2019).

Observational evidence from the Lower Keys surveys suggested that there could have been mangrove-coral habitats with higher coral diversity on some of the more open-water shorelines but that they were destroyed, possibly during the passage of Hurricane Irma, which made direct landfall as a category 4 storm on Cudjoe Key in September 2017. Coral rubble from multiple species was observed in uncompacted sediment layers among mangrove prop roots at both oceanside (east of Cook Island) and backcountry (Johnston Key mangroves) sites. It is possible that the coral rubble was transported to these sites by the storm. However, dead coral nubbins that remained attached to the substrata could be felt beneath the sediment layer along the mangrove fringe at the Cook Island site. In the backcountry, there were several sites along the Gulf of Mexico-facing shore where the mangrove prop roots had been scoured clean (e.g., Johnston Key Mangroves and Sawyer Key). Sawyer Key had up to 1-m thick wrackline of seagrass and sponges along the shore and the Snipe Keys had a layer of storm mud in the mangroves. Although the hardbottom extended all the way to the mangrove shoreline in many of these areas, there was a layer of unconsolidated sediment 5 to 15 cm thick covering it, impeding coral survival close to the mangroves. These observations are consistent with reports of storm damage in the mangroves after Hurricane Irma. Radabaugh et al. (2019) reported widespread mortality in Lower Keys mangroves and sedimentary storm-surge deposits ranging from 1–7 cm thick. Additionally, severe shoreline erosion occurred in several locations and seagrass wrack along some mangrove shorelines was 5–15 cm thick in the months immediately after the storm (R Moyer, 2017, unpublished data). These open-water shorelines appear to be prime potential coral habitat (clear, oceanic water combined with hardbottom and mangrove-lined shoreline). From the observed coral rubble, scrubbed prop roots, and unconsolidated sediment layer, we infer that there may have been prop-root- or channel-coral habitat in these areas, but that Hurricane Irma destroyed them. This type of destruction in the highly diverse mangrove-coral habitat in the U.S. Virgin Islands was documented in the wake of Hurricanes Irma and Maria in 2017 (Rogers 2019). This suggests that these areas may be worth reassessing in 3 to 5 years to see if new diverse coral communities become established as the mangrove habitats continue to recover.

Due to time, weather, and funding limitations, our surveys did not include all possible mangrove shoreline targets in the Florida Keys, so additional locations with mangrove-coral habitats are likely yet to be identified. There are over 1,400 linear km of mangrove shoreline in the Lower Keys alone [estimated from http://geodata.myfwc.com/datasets/esi-shoreline-classification-lines-florida]. While the survey approach employed in this study used informed decisions to target those areas with the highest probability of hosting mangrove-coral habitats, some areas that were missed by this initial effort may host even higher coral diversity than the ones documented here. Over 30 species of scleractinian corals have been described in mangrove habitats of the U.S. Virgin Islands, demonstrating that mangroves can host a high-diversity assemblage of corals if the environmental conditions are favorable (Rogers 2017).

## Conclusions

This study was a first effort to locate and characterize mangrove-coral habitats in the Florida Keys. We documented areas where corals were growing directly on and under mangrove prop roots (prop-root-coral habitats) and where they were growing under the shade of the mangrove canopy (channel-coral habitats). Areas with corals growing on prop roots were characterized by roots hanging into undercut channels and/or with strong tidal currents and often connections to adjacent open-ocean waters. Coral species found growing on and directly adjacent to prop roots included *P. porites* (multiple morphs), *S. radians* and *F. fragum*. Channel-coral habitats predominantly hosted *S. radians*, although single colonies of *Solenastrea bournoni* and *Stephanocoenia intersepta* and several *S. siderastrea* were observed. There is circumstantial evidence that suggests additional mangrove-coral habitats existed on oceanside and backcountry islands but were destroyed by Hurricane Irma. These mangrove-coral habitats may be refugia for corals threatened by climate change and disease outbreaks. Further evaluation is needed to determine if these habitats could contribute to coral restoration efforts; for example, as a source of adapted high-resilience genetic materials, or as locations to support the growth and acclimation of coral outplants in areas that may be at lower risk of coral bleaching, disease, or storm damage.

## Acknowledgements

The authors thank J. Voelschow for assisting with data formatting, B. Williams for converting KMZ files to ArcGIS maps and estimating the area surveyed via track lines, and B. Boynton for layout of photo plates. We are grateful to A. Franklin for geospatial technology support. Any use of trade, firm, or product names is for descriptive purposes only and does not imply endorsement by the U.S. Government or the State of Florida.

